# Transcriptional regulation profiling reveals disrupted lipid metabolism in failing hearts with a pathogenic phospholamban mutation

**DOI:** 10.1101/2020.11.30.402792

**Authors:** Jiayi Pei, Renee G.C. Maas, Ema Nagyova, Johannes M.I.H. Gho, Christian Snijders Blok, Iris van Adrichem, Jorg J.A. Calis, René van Es, Shahrzad Sepehrkhouy, Dries Feyen, Noortje van den Dungen, Nico Lansu, Nicolaas de Jonge, Hester M. den Ruijter, Manon M.H. Huibers, Roel A. de Weger, Linda W. van Laake, Marianne C. Verhaar, Peter van Tintelen, Frank G. van Steenbeek, Alain van Mil, Jan W. Buikema, Boudewijn Burgering, Ioannis Kararikes, Mark Mercola, Pieter A. Doevendans, Joost Sluijter, Aryan Vink, Caroline Cheng, Michal Mokry, Folkert W. Asselbergs, Magdalena Harakalova

**Affiliations:** Department of Cardiology, Division Heart & Lungs, University Medical Center Utrecht, University of Utrecht, Utrecht, The Netherlands; Regenerative Medicine Center Utrecht, University Medical Center Utrecht, Utrecht, The Netherlands; Department of Nephrology and Hypertension, Division of Internal Medicine and Dermatology, University Medical Center Utrecht (UMCU), The Netherlands; Epigenomics facility, University Medical Center Utrecht, Utrecht, The Netherlands; Department of Pathology, University Medical Center Utrecht (UMCU), Utrecht, The Netherlands; Stanford Cardiovascular Institute and Department of Medicine, Stanford University, Stanford, CA, USA; Department of Genetics, University Medical Center Utrecht, Utrecht, The Netherlands; Department of Clinical Sciences, Faculty of Veterinary Medicine, University of Utrecht, NL; Center for Molecular Medicine, Molecular Cancer Research Section, University Medical Center Utrecht, The Netherlands; Netherlands Heart Institute, Utrecht, The Netherlands; Wilhelmina Children’s Hospital, Department of Pediatric Gastroenterology, Division Child Health, Utrecht, the Netherlands; Institute of Cardiovascular Science, Faculty of Population Health Sciences, University College London, London, United Kingdom; Health Data Research UK and Institute of Health Informatics, University College London, London, UK

**Keywords:** phospholamban, heart failure, DNA regulatory region, chromatin regulation, H3K27ac, transcription factor binding motif

## Abstract

**Background:** The R14del mutation in the phospholamban (*PLN*) gene is associated with various types of cardiomyopathies and increases the risk of developing life-threatening ventricular arrhythmias. In this study, we focused on a homogeneous Dutch founder cohort of genetic cardiomyopathy due to PLN R14del mutation and aimed to study the influence of epigenetic changes from a multi-dimensional perspective.

**Results:** Using cardiac tissue of PLN R14del patients and donors, we identified differentially acetylated promoters and enhancers (H3K27ac ChIPseq), annotated enriched transcription factor (TF) binding motifs located in those regions, and identified differentially expressed genes (RNA-seq). In line with the fibrofatty replacement in PLN R14del hearts at the histological level, our integrative analysis detected the downregulation of key TF regulators in fatty acid oxidation (FAO) metabolisms and their downstream target in PLN R14del hearts as compared to controls. We further examined heart tissue using immunofluorescence staining (IF) and to confirm the mitochondrial lipid abnormalities in the PLN R14del hearts. Furthermore, we observed the accumulation and deformation of lipid droplets and a disrupted morphology of mitochondria, the key organelle where FAO takes place, in PLN R14del heart using transmission electron microscopy (TEM).

**Conclusion:** Using multi-omics approaches, we successfully obtained a unique list of chromatin regions and genes, including TF-coding genes, which played important roles in the metabolism-related signalling in PLN R14del hearts.

## INTRODUCTION

The R14del mutation in the phospholamban (*PLN*) gene is associated with dilated cardiomyopathy (DCM) and/or arrhythmogenic (biventricular) cardiomyopathy (ACM) with a high risk for life-threatening ventricular arrhythmias.^1,2^ Unfortunately, no effective treatment is currently available for PLN R14del patients. This mutation explains a large proportion of Dutch DCM and ACM cases, 15% and 12%, respectively, which is linked to a shared haplotype originated around the year 1400 A.D. in the northern parts of The Netherlands.^3,4^ Additional families carrying the PLN R14del mutation have been detected in other European countries, the United States, and Canada.^5^ PLN is a small phosphoprotein located in the cardiomyocyte sarcoplasmic reticulum (SR) acting as the major regulator of SERCA2a/ATP2A2 activity and calcium (Ca2+) cycling. Previous research in PLN R14del cardiomyocytes confirmed disturbed Ca2+ cycling and abnormal cytoplasmic distribution of the PLN protein.^1,6^ Aggregation, aggresome formation, and autophagy of the mutant PLN R14del protein in patient cardiac tissue have also been described.^7^ Macroscopically, the end-stage PLN R14del heart shows a biventricular subepicardial fibrofatty tissue replacement of the myocardium, which is accompanied by profound interstitial fibrosis, transmural adipocyte infiltration, and islands of isolated cardiomyocytes between the adipocytes.^8–10^

Adipocyte infiltration and fibrosis in the myocardium serve as an anatomical barrier and thus create a substrate for fatal arrhythmias resulting from re-entry of the electrical signal.^11^ The distribution of fibrofatty tissue replacement in PLN R14del carriers measured by late gadolinium enhancement can distinguish mutation carriers with and without ventricular arrhythmias.^12^ Many diseases, such as the various types of acquired (e.g. ischemic) and genetic (e.g. due to mutations in *PKP2, DSG2, DSP, DSC2, JUP, SCN5A*, and *TNNT2)* cardiomyopathies, myotonic dystrophy, obesity, and atrial fibrillation, exhibit fibrofatty infiltration as one of the pathophysiological hallmarks.^11,13–15^ However, it is still unclear which cell type(s) or mechanisms are responsible for the adipocyte infiltration by either the activation of the already existing pool of adipocytes or transdifferentiation of (cardiac) cells into adipocytes.^11^ Interestingly, next to the extracellular adipocyte infiltration, abnormalities in intracellular cardiomyocyte lipid metabolism have been observed in arrhythmogenic right ventricular cardiomyopathy (ARVC) caused by mutations in *PKP2*. The production of lipid droplets and adipogenic markers in *PKP2* mutant cardiomyocytes have been observed.^16^ Induced pluripotent stem cell-derived cardiomyocytes (iPSC-CMs) with a homozygous *PKP2* mutation showed aggressive lipogenesis, elevated apoptosis, and reduced metabolic flexibility.^17^ Furthermore, there are several forms of childhood cardiomyopathies caused by mutations in the mitochondrial fatty acid oxidation (FAO) pathway genes, such as *HADHA, HADHB, CPT2, ACADVL*, which have shown intracellular cardiomyocyte lipid droplet storage and adipocyte infiltration in the myocardium and other organs. Given the fact that lipid droplets and adipocyte infiltration is observed in various types of (genetic) forms of cardiac remodelling, a metabolism-related mechanism could be responsible for the progression of cardiomyopathies in general.

During heart failure, stressed cardiomyocytes suffer from an impaired capacity to oxidize fatty acids.^18^ The failing heart subsequently switches to glycolysis for energy production,^19^ resembling the energy balance of the fetal heart. However, the compensatory increase of glucose oxidation in the fetal-like energy management manner is not sufficient to meet adult energy consumption needs and will disturb the capacity of FAO-based ATP synthesis, leading to further starvation of the heart.^20,21^ Two key regulators of glucose and fatty acid metabolism on a molecular level, glucose transporter 1 (*GLUT1*) and peroxisome proliferator-activated receptor alpha (*PPARA*), respectively, are both downregulated in failing hearts,^22^ indicating the insufficient energy production. Accumulation of lipid droplets has been shown in murine hearts by suppressing *PPARA* expression.^23^ Myocardial adipogenesis in patients with cardiomyopathies, including PLN R14del carriers, is considered as an aberrant remodelling that is associated with the myocardial loss.^24^ Nevertheless, the metabolic adaptation and the precise pathophysiological mechanisms in PLN R14del mutation carriers involved in severe fat infiltration remains unknown.

Here, we employed multiple next-generation-sequencing approaches in a hypothesis-free manner to study the compact myocardial area of PLN R14del patients and healthy control hearts, in which the fatty epicardial layers were removed. We first profiled the activity of H3K27ac assayed by chromatin immunoprecipitation and sequencing (ChIP-seq) in human cardiac tissue from PLN R14del cardiomyopathy patients in comparison to healthy donor hearts. Based on the list of differentially acetylated regulatory regions between PLN patients and controls, we performed *in silico* region-to-gene annotations and TFBMs enrichment analysis. We also obtained the transcriptional changes in PLN versus control hearts using RNA sequencing (RNA-seq). By integrating the information at the DNA and RNA levels, we identified a set of TFs, which played a critical role in the end-stage PLN R14del hearts, including the potential upstream regulators of the energy metabolism.

## METHODS

### Study design and samples

This study was approved by the Biobank Research Ethics Committee, University Medical Center Utrecht, Utrecht, The Netherlands, under protocol number 12-387 (cardiac tissues). Written informed consent was obtained or in certain cases waived by the ethics committee when obtaining informed consent was not possible due to the death of the individual. Heart samples collected at autopsy or transplantation were obtained from a homogeneous cohort of patients who all carried the same pathogenic PLN R14del mutation (n=6). Four control hearts obtained from unused organ donors (n=3) or from autopsy (n=1) were used as a reference. To further elaborate on PLN R14del-specific changes, hearts from patients with ischemic cardiomyopathy (n=4) and from non-ischemic cardiomyopathy based on mutations in genes encoding sarcomeric proteins (n=6) were also included. An overview of cardiac tissues is presented in **Table S1**.

### Cardiac tissue handling

Cardiac tissues used for ChIP-seq and RNA-seq were obtained from regions halfway between the atrioventricular valves and the apex and were stored at −80°C. From each individual, an adjacent block of tissue from the same biopsy was used for ChIP-seq was paraffin-embedded and stained with Masson’s trichrome. If applicable, macroscopically visible regions of subepicardial fat or myocardial fibrofatty replacement were removed from frozen tissue blocks and 12-30 tissue slices of 10 μm thick were cut to achieve a comparable amount of myocardial tissue in the sequenced material. For all the other samples ten 10 μm thick frozen slices were collected.

### Chromatin H3K27ac immunoprecipitation and sequencing of human cardiac tissues

Chromatin was isolated from frozen cardiac tissues using the MAGnify™ Chromatin Immunoprecipitation System kit (Life Technologies) according to the manufacturer’s instructions. In brief, the obtained cardiac tissue was crosslinked with 1% formaldehyde and the crosslinking was stopped by adding 1.25 M glycine. Cells were lysed using the kit-provided lysis buffer and nuclei were sonicated using Covaris microTUBE (duty cycle 5%, intensity 2, 200 cycles per burst, 60s cycle time, 10 cycles). Sheared chromatin was diluted based on the expected number of isolated cells and was incubated with an anti-H3K27ac antibody (ab4729, Abcam) pre-coupled to magnetic beads for 2 hours at 4°C. Beads were extensively washed and crosslinking was reversed by the kit-provided reverse crosslinking buffer with 20 mg/mL Proteinase K. DNA was purified using ChIP DNA Clean & Concentrator kit (Zymo Research). Isolated DNA was additionally sheared, end-repaired, sequencing adaptors were ligated and the library was amplified by PCR using primers with sample-specific barcodes according to our modification to manufacturer’s recommendations. After PCR, the library was purified and checked for the proper size range and for the absence of adaptor dimers on a 2% agarose gel and sequenced on SOLiD Wildfire sequencer.

### H3K27ac ChIP-seq analyses

Sequencing reads were mapped against the reference genome (hg19 assembly, GRCh37) using the BWA package (−c, −l 25, −k 2, −n 10).^25^ Multiple reads mapping to the same location and strand have been collapsed to single reads and only uniquely placed reads were used for peak/region calling. Regions were called using Cisgenome 2.0 (−e 150 -maxgap 200 –minlen 200).^26^ Next, to obtain a common reference, region coordinates from all PLN and control samples were stretched to at least 2000 base pairs and collapsed into a single common list. Overlapping regions were merged based on their outmost coordinates. Only the autosomal regions supported by at least 2 independent datasets were further analyzed. Sequencing reads from each ChIP-seq library were overlapped with the common region list, to set the H3K27ac occupancy for every region-sample pair. Obtained regions were further examined in 4 analyses (**Table S2**).

#### Detection and gene annotation of differentially acetylated regions (Analysis 1)

Regions with differential H3K27ac occupancy between PLN and control hearts were identified using DESeq2 standard settings (p<0.05 as calculated by Wald test)^27^ and are referred to as ‘differentially acetylated regions’. Supervised hierarchical clustering was performed with quantile normalized (limma::normalizeQiantiles() function in R), log2 transformed and median centred read counts per common region. To avoid the log2 transformation of zero values, one read was added to each region. Region to gene annotation was performed in silico using a conservative window of +/−5kb from the transcription start site (TSS).

#### Enrichment of TF binding motifs (TFBM) in differentially acetylated regions (Analysis 2)

A total of 3396 cardiac DNAse hypersensitivity sites (DHS) obtained from the ENCODE database (Heart_OC, Primary frozen heart tissue from NICHD donor ID:1104, Male, Caucasian, 35 years old)^28^ overlapping with differentially acetylated regions in the PLN vs. control group (both hypo- and hyperacetylated) were used for this analysis. The genomic sequence of DHS was repeat-masked and the enrichment of TFBM was calculated against the shuffled sequences using the Analysis Motif of Enrichment (AME tool) of the MEME Suite^29^ with the following settings: motif database: human (HOCOMOCO v9), background model sequence set to 0.29182,0.20818,0.20818,0.29182, pseudo count added to a motif column: 0.25, Wilcoxon rank-sum test (quick), p<0.05), number of multiple tests for Bonferroni correction: #Motifs× #PartitionsTested = 426×1 = 426. The functional annotation of the enriched TFs was performed as described above.

#### PLN-specificity analysis of differentially acetylated regions (Analysis 3)

Sequencing reads from each ChIP-seq sample (PLN (n=6), control (n=4), ischemic (n=4), and sarcomeric (n=6)) were compared to the common differentially acetylated region list to set the H3K27ac occupancy for every region-sample pair. Raw read counts were quantile normalized (limma::normalizeQiantiles() function in R), log2 transformed and median centred (to avoid log2 transformation of zero values, one read was added to each region). The median value from each sample group was used to construct an n x× k table where n = 4 (one value per each sample type) and k represent the number of differentially acetylated regions. The k-means (nstart = 200) function in R was used to partition the regions into 12 different clusters. To enable the reproducibility of the identified clusters set.seed(10) R command was called before the clustering. Annotation of clusters with PLN-specific patterns was performed as described above.

#### Pathway analysis of genes annotated to differentially acetylated regions (Analysis 4)

ToppFun and STRING were used for gene list enrichment analysis and candidate gene prioritization based on functional annotations and protein interaction networks.^30,31^ For ToppFun, the list of hyper- and hypoacetylated genes was tested using probability density function p-value calculation method, FDR B&H correction, p-value cut-off of 0.05, and gene limit of 1-2,000 genes per term. Since the pre-build gene/protein networks integrated into ToppFun were not created using the same criteria and the annotated number of genes varies significantly, we reported also genes belonging to known disease pathways even below the p-value threshold (where indicated). For protein network interaction visualization STRING v10.0 was used with a minimum required interaction score at the highest confidence setting for all differentially acetylated peaks.

### Transcriptome analysis of human cardiac tissues using RNA sequencing

RNA was isolated using ISOLATE II RNA Mini Kit (Bioline) according to the manufacturers’ instructions with minor adjustments. After the selection of mRNA, libraries were prepared using the NEXTflexTM Rapid RNA-seq Kit (Bioo Scientific). Libraries were sequenced on the Nextseq500 platform (Illumina), producing single-end reads of 75bp. Reads were aligned to the human reference genome GRCh37 using STAR v2.4.2a.^32^ Picard’s AddOrReplaceReadGroups v1.98 (http://broadinstitute.github.io/picard/) was used to add read groups to the BAM files, which were sorted with Sambamba v0.4.5 and transcript abundances were quantified with HTSeq-count v0.6.1p1 using the union mode.^33,34^ Subsequently, reads per kilobase per million mapped reads (RPKMs) were calculated with edgeR’s RPKM function.^35^ DESeq2 was used to identify differentially expressed regions using the cutoff of p<0.05 in the Galaxy environment (default settings).^36^

### Availability of data and materials

All relevant data are available within the article and the supplementary files. Because of the sensitive nature of the data collected for this study, requests to access additional dataset from qualified researchers trained in human subject confidentiality protocols may be sent to the corresponding authors.

## RESULTS

### Detection of differentially acetylated regions in PLN versus control hearts

Using ChIP-seq on cardiac tissues we identified 28,149±9,538, and 25,721±8,460 regions with H3K27ac binding in PLN and control hearts, respectively (**Table S2**). Next, we combined regions that were identified in at least two independent samples into a set of 23,356 regions to assess differentially acetylated regions between two groups. In total, 2,107 autosomal regions showed differential H3K27ac levels (**Figure 1A**). Among these, 958 and 1,149 regions were identified as hyper- and hypoacetylated in PLN hearts as compared to the controls, respectively (**Table S3A**). Thus, chromatin isolated from end-stage heart failure tissue of explanted hearts provided a reasonable number of differentially acetylated regions for further annotation.

**Figure 1:**
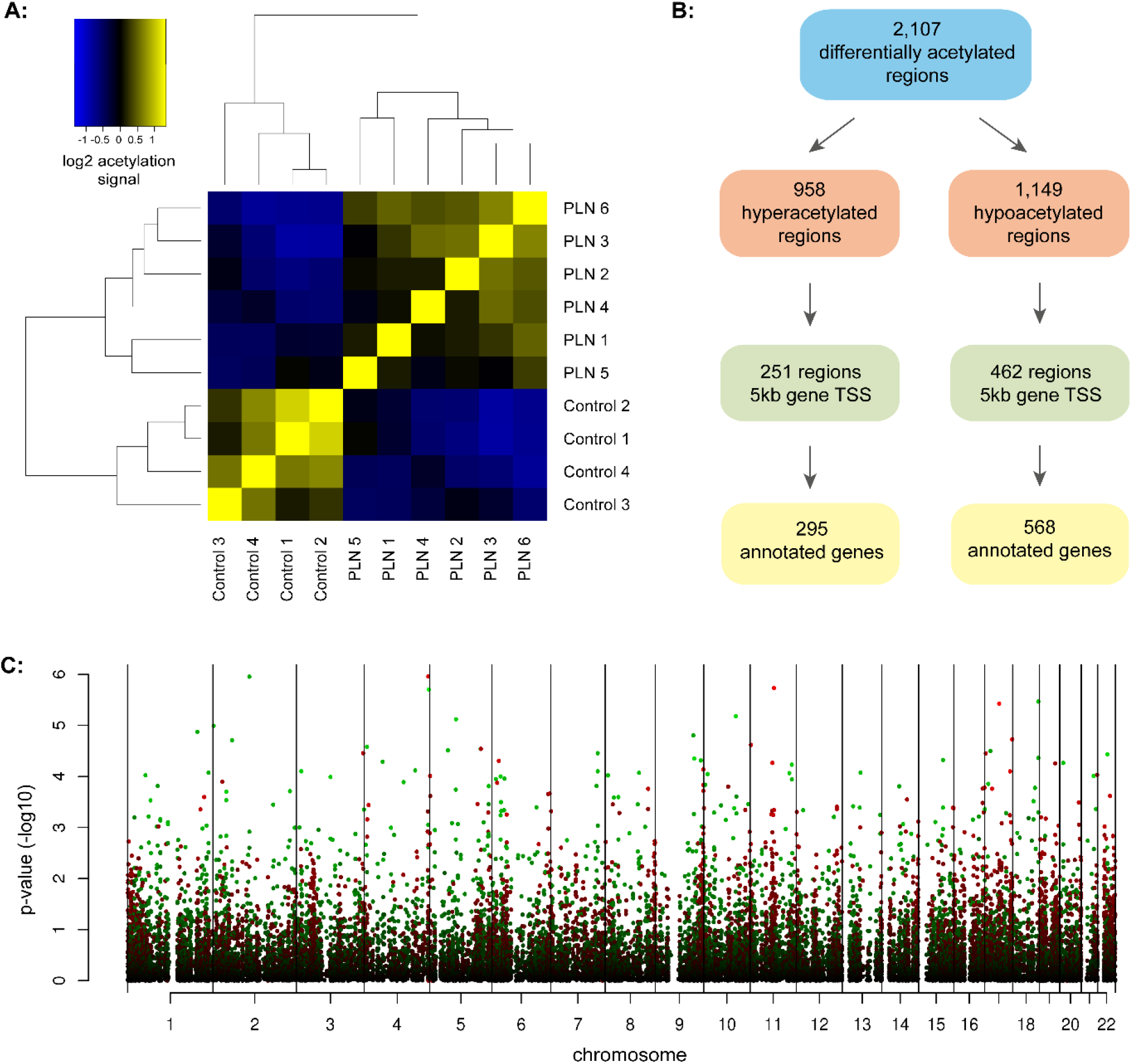
Detection and gene annotation of differentially acetylated regions (ChIP-seq): **A:** Correlation heat map of all differentially acetylated regions in cardiac tissue from PLN patients vs. controls. The supervised clustering is based on the mean of H3K27ac signal over differentially acetylated regions in the heart (Wald test p-value <0.05) and separates the group of PLN patients and controls. **B:** Flowchart representing the detection and annotation of differentially acetylated regions in PLN patients vs. controls. Hyperacetylation indicated signal higher in PLN group and hypoacetylation signal lower in PLN group as compared to controls. **C:** Manhattan plot representing the distribution of tandem regulated chromatin domains detected in PLN and control group across autosomal chromosomes. Color of dots represents differences in region H3K27 acetylation: non-significant regions (black), hyperacetylated regions (green) and hypoacetylated regions (red).

### Gene annotation of differentially acetylated regions

Next, we focused on differentially acetylated peaks in close vicinity to gene bodies. We have chosen a conservative cutoff and followed only genes with differentially acetylated regions using a window of ±5kb between the peak centre and transcription start site (TSS) of a gene. In total, 295 genes were identified in the close vicinity to 251 hyperacetylated peaks, including 233 protein-coding genes, 2 pseudogenes, 38 long and 22 short non-coding RNA genes. We have also detected 568 genes close to 462 hypoacetylated peaks, including 476 protein-coding genes, 5 pseudogenes, 53 long and 34 short non-coding RNA genes (**Figure 1B**, **Table S3B, and 3C**). Tandem regulated chromatin domains based on the distribution of differentially H3K27 acetylated regions in patients vs. controls are shown in **Figure 1C**. Selected examples of regions in the vicinity of 4 cardiac genes (*FST, SMAD7, PROB1,* and *MYO15B*) compared to H3K27ac signals from the ENCODE project are shown in **Figure 2A**.

**Figure 2:**
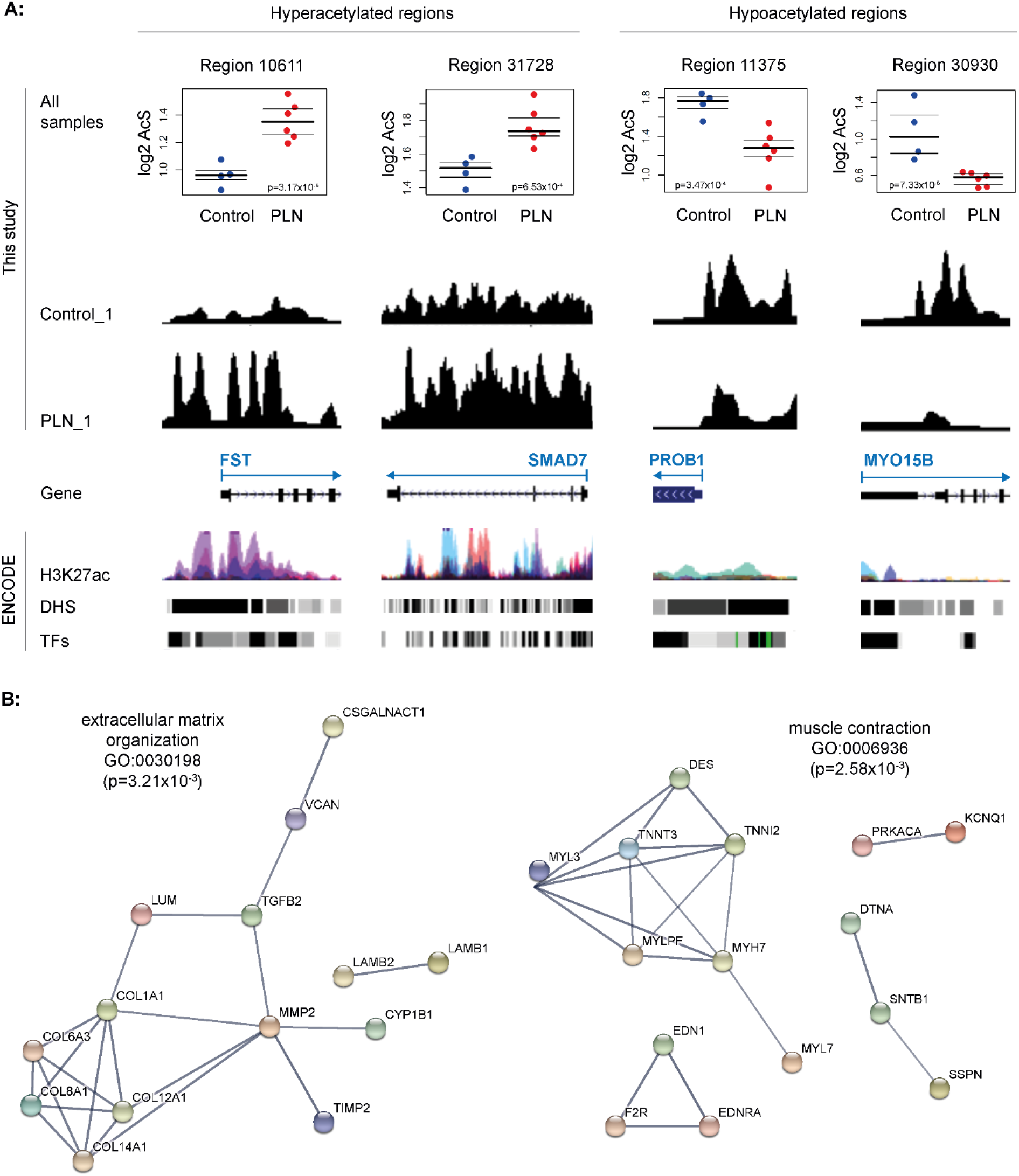
Selected examples of differentially acetylated regions (ChIP-seq): **A:** Four differentially acetylated regions depicted using the UCSC Genome Browser. In each example (*FST, SMAD7, PROB1,* and *MYO15B*) the region overlaps with the putative promoter (upstream region, 5’UTR, first exons in region tracks) and shows a significant difference between PLN patients and controls (dot plots). Arrow beginning indicates gene transcription start site (TSS). AcS = acetylation signal. ENCODE = publicly available ENCODE consortium data default display. H3K27ac = layered H3K27ac ChIPseq data in 6 cell types (GM12878 - red, H1-hESC – orange, HSMM – green, HUVEC – light blue, K562 – dark blue, NHEK – purple, NHLF – pink). DHS – DNaseI hypersensitivity clusters in 125 cells, TFs = ChIPseq for 161 TFs. More details about the visualized H3K27ac ChIPseq library tracks (PLN_1 and Control_1) are shown in Table S1 and S2. **B:** Selected examples of STRING protein database used to detect interacting proteins annotated to differentially annotated regions. Input list contained 863 genes annotated to regions’ vicinity. Only the highest confidence interactions are displayed. Disconnected nodes were removed from the network image. Displayed genes belong to the indicated GO terms (extracellular matrix organization and muscle contraction).

### Pathway analysis of genes annotated to differentially acetylated regions

To predict altered biological processes in PLN hearts when compared with controls, we performed gene enrichment analysis of 863 genes annotated to differentially acetylated regions using ToppFun (padj<0.05). The analysis of 295 hyperacetylated genes resulted in several enriched GO terms and pathways of various sources related to three main groups: 1. fibrosis (extracellular matrix components, collagen synthesis, TGF-beta signalling pathway, cell adhesion, immune system), 2. (cardiovascular) development and 3. chromatin assembly (histone function, nucleosome, cell senescence and proliferation, **Table S4A**). Significantly enriched processes related to chromatin assembly were mostly based on the core histone H1 cluster on chromosome 6 based on 5 neighbouring significantly hyperacetylated regions. On the other hand, gene enrichment analysis of hypoacetylated 568 genes resulted in two main groups: 1. (lipid) metabolism (fatty acid beta-oxidation, transferase activity) and 2. mitochondrial function (**Table S4B**). Examples of two Gene Ontology (GO) processes relevant to cardiomyopathy development are depicted in **Figure 2B** (extracellular matric organization and muscle contraction).

### Enrichment of TF binding motifs in differentially acetylated regions

Next, we studied TFBMs that were enriched in the differentially acetylated regions, which might be involved in the pathogenesis of the end-stage heart failure. We employed the AME program to test the enrichment of TFBMs using the DNA sequence of differentially acetylated peaks and detected enrichment in 202 TFBMs and annotated them to 200 TF-encoding genes (**Table S5A**). In line with previous results, gene enrichment analysis of annotated TFs pointed to similar biological processes, among others, those involved in adipogenesis, myogenesis, mitochondrial structure, TGF beta signalling in fibrosis, heart development, and cell death (**Table S5B**). Protein interactions of several TFs linked to metabolic pathways such as adipogenesis, mitochondrial organization, and of lipid metabolism by PPARA, are shown in **Figure 3A**.^37^ Interestingly, the regulatory regions in the close vicinity of gene bodies encoding seven TFs with enriched TFBM also showed differential acetylation level: *ESR1, ESRRA, KLF3, MAFG, MYC, NFATC2,* and *ZFHX3*. We confirm that enriched TFBMs inside differentially acetylated regions point to TFs working together with genes with differentially acetylated regulatory regions in the development of heart failure. Examples of several TFs with their enriched TFBMs are shown in **Figure 3B**.

**Figure 3:**
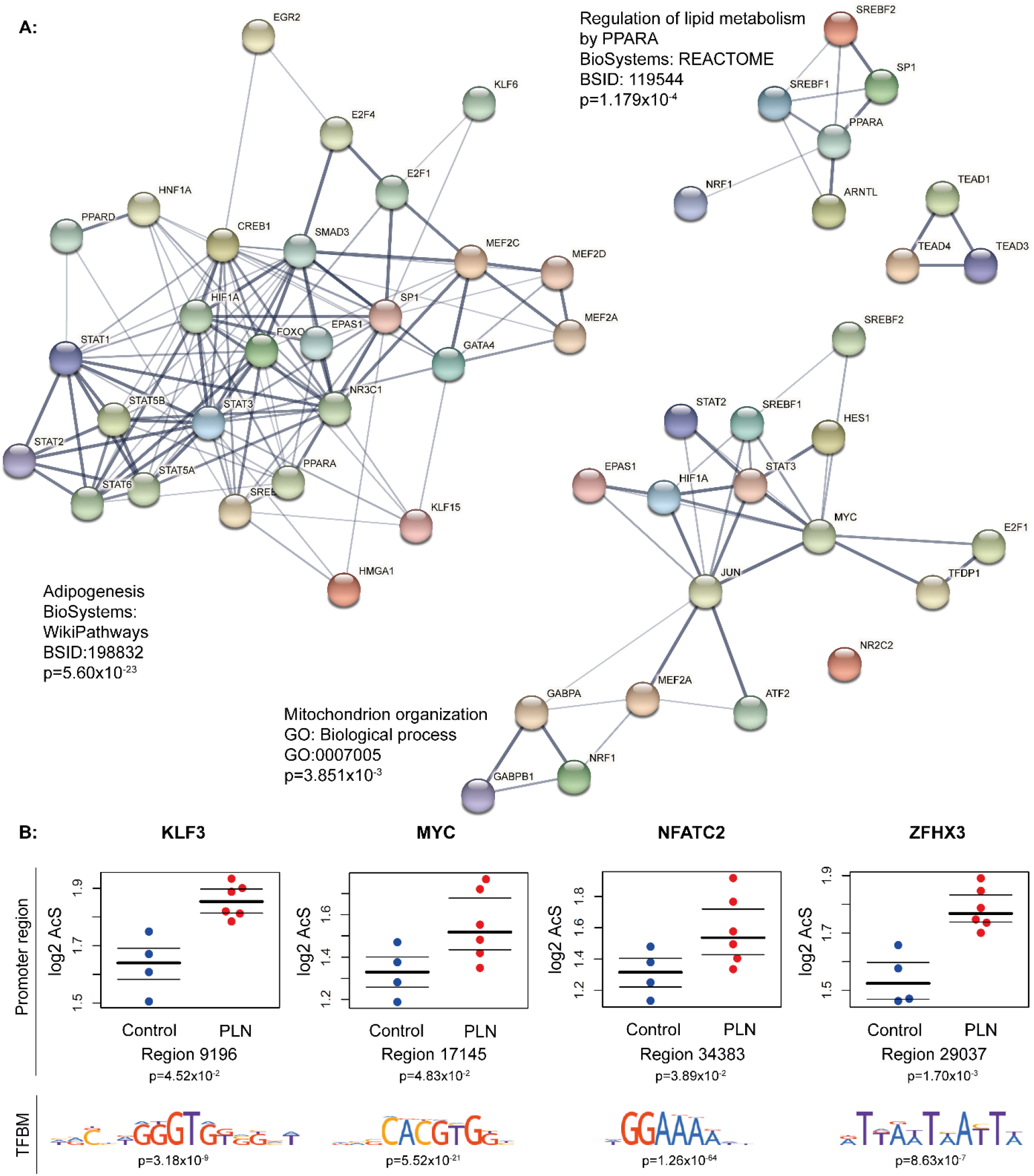
Enrichment of TF binding motifs in differentially acetylated regions (ChIP-seq): **A:** Selected examples related to mitochondrial (lipid)metabolism: Adipogenesis (Wiki Pathways - containing *PPARA* and *KLF15*), Mitochondrion organization (GO), and Regulation of lipid metabolism by PPARA (REACTOME) using STRING protein database to detect interacting TFs annotated by both hyper- and hypoacetylated regions. Only the highest confidence interactions are displayed. **B:** Selected examples of TFs with both differentially acetylated promoter as well as enrichment of binding motif (source: HOCOMOCO) in the differentially acetylated regions (*KLF3, MYC, NFATC2, and ZFHX3*). Dot plots represent the acetylation signal (AcS) measured for each sample.

### PLN-specificity analysis of differentially acetylated regions

In order to further characterize specific pathways involved in PLN R14del and to exclude general events linked to end-stage heart failure, we also compared H3K27ac profiles of patients with other types of ischemic cardiomyopathy (n=4) and non-ischemic dilated cardiomyopathy (n=6). K-mean analysis revealed PLN R14del-specific clusters of regions, among others, with PLN-specific hyper- and hypoacetylation or clusters of regions linked to cardiomyopathies in general (**Table S6A**, **Figure 4A**). Next, we focused on the main clusters showing a PLN R14del-specific pattern. We have identified two clusters with hyperacetylation (clusters 1 and 12) projecting to 50 genes and two clusters with hypoacetylation (clusters 3 and 4) resulting in 158 genes in PLN 14del when compared with the other groups (**Table S6B**). We show PLN R14del-specific patterns of eight regions in the vicinity of genes, such as *LINC01140, WIPF1, OSR1, IFFO2, LTBP1, GCGR*, or *CNNM4* (**Figure 4B**). Interestingly, at least 13 genes from the hyperacetylated group and 55 genes from the hypoacetylated group are known to be involved in various pathways related to (phospho)lipid and lipoprotein synthesis and metabolism, such as recognizable genes *DGKZ, HADHA/HADHB, INPP5J, MLYCD, PLCD3,* and *PLPP1* **(Figure 5)**. Here, by comparing differentially acetylated regions in PLN patients to other types of dilated cardiomyopathy, we detected several PLN-specific regions.

**Figure 4:**
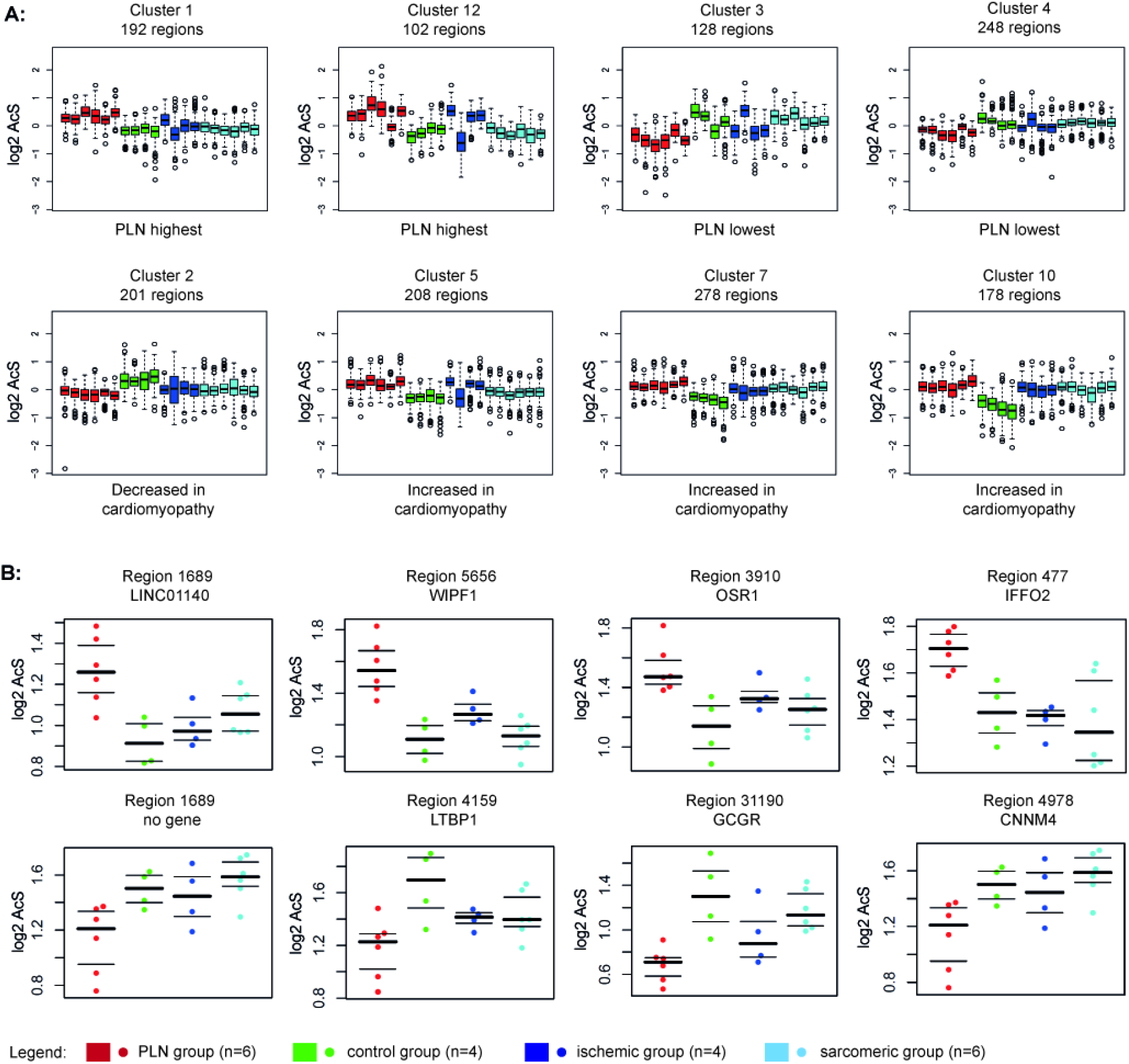
PLN-specificity analysis of differentially acetylated regions (ChIP-seq): K-mean clustering of PLN R14del cardiomyopathy (red), healthy controls (green), ischemic cardiomyopathy (dark blue) and sarcomeric non-ischemic cardiomyopathy (light blue) based on H3K27ac signal was used to partition the regions into 12 different clusters. **A)** K-mean clusters showing PLN-specific hyper- and hypoacetylation regions when compared with controls and hearts with ischemic or non-ischemic dilated cardiomyopathy in general; **B)** Examples of 8 PLN-specific regions and genes in the vicinity of these regions (*LINC01140, WIPF1, OSR1, IFFO2, LTBP1, GCGR*, or *CNNM4)*. No genes have been annotated in the vicinity of region 1689.

**Figure 5:**
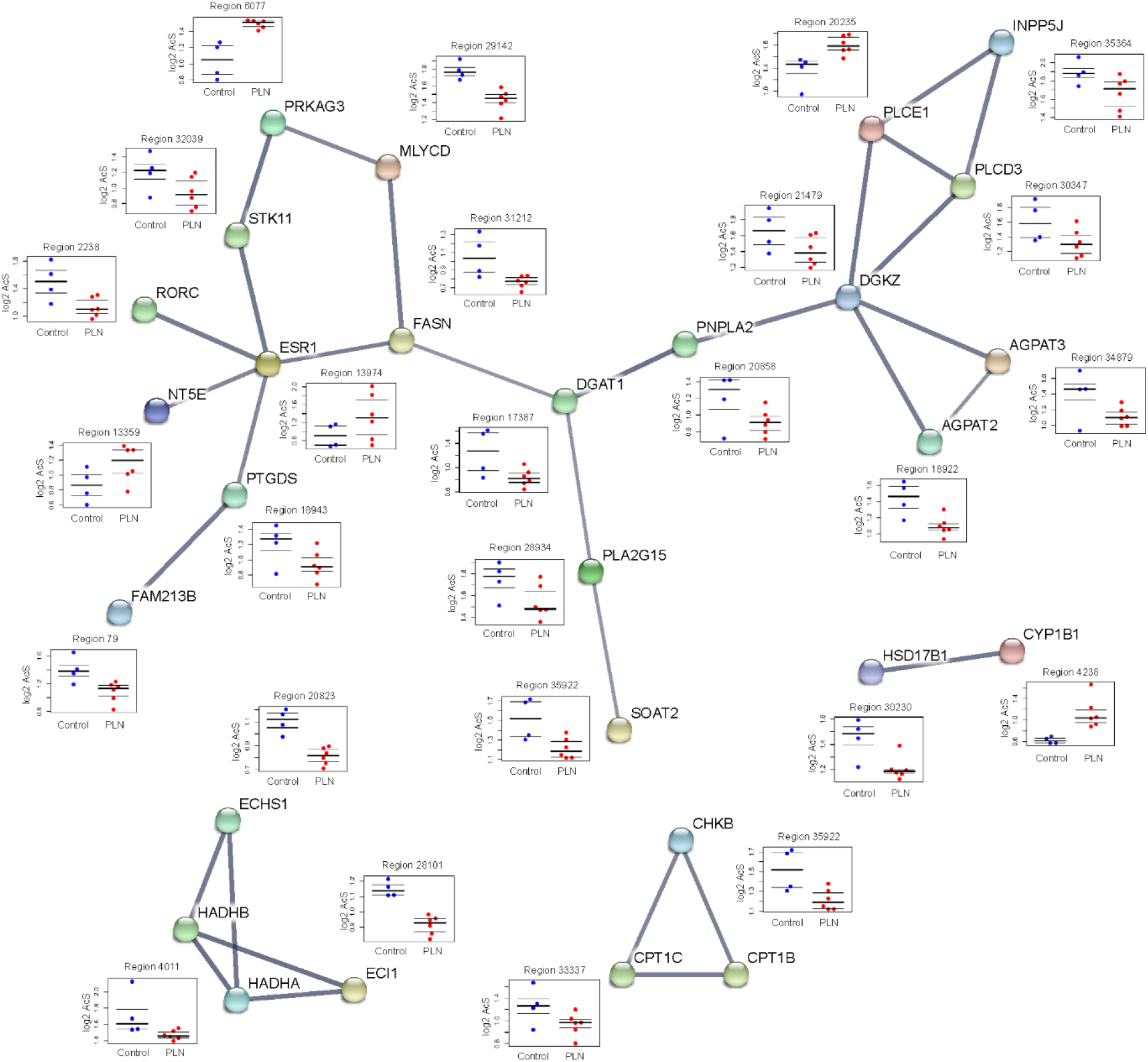
Genes annotated to PLN-specific differentially acetylated regions (ChIP-seq): As an effort to identify differentially acetylated regions in the vicinity of genes involved in lipid metabolism possibly linked to fibrofatty tissue replacement in PLN group, this figure shows selected examples of STRING protein database used to detect interacting proteins annotated to differentially annotated regions as part of the GO term ‘lipid metabolic process’ (GO:0006629). Only the highest confidence interactions are displayed. Disconnected nodes were removed from the network image. Dot plots represent the acetylation signal (AcS) measured for each sample.

### Detection of differentially expressed genes in PLN versus control hearts

Using RNA-seq on cardiac tissues from the exactly same PLN and control hearts, we obtained 3,541 differentially expressed genes between PLN and control hearts. Among these, 1,668 and 1,873 genes were identified as up- and downregulated genes in the PLN group as compared with the control group, respectively (**Table S7**). Like the isolated chromatin, mRNAs isolated from end-stage heart failure tissue of explanted hearts also provided a reasonable number of differentially expressed genes for further annotation.

### Pathway analysis of differentially expressed genes

To study the biological processes that occur in PLN hearts as compared with the controls, we examined the pathways enriched by differentially expressed genes. Consistent with the results using annotated genes from differentially acetylated regions, enriched GO terms and pathways by 1,668 upregulated genes were mostly involved in fibrosis (extracellular matrix organization, cell adhesion, collagen binding, TGF-beta signalling pathway), and cardiovascular development (circulatory system development, blood vessel development, **Figure 6AB**, **Table S8A**). Gene set enrichment analysis of 1,873 downregulated genes was dominantly involved in the energy metabolism (oxidative phosphorylation, ATP synthesis coupled proton transport, acetyl-CoA C-acyltransferase activity) and the mitochondrial function (**Table S8B**).

**Figure 6:**
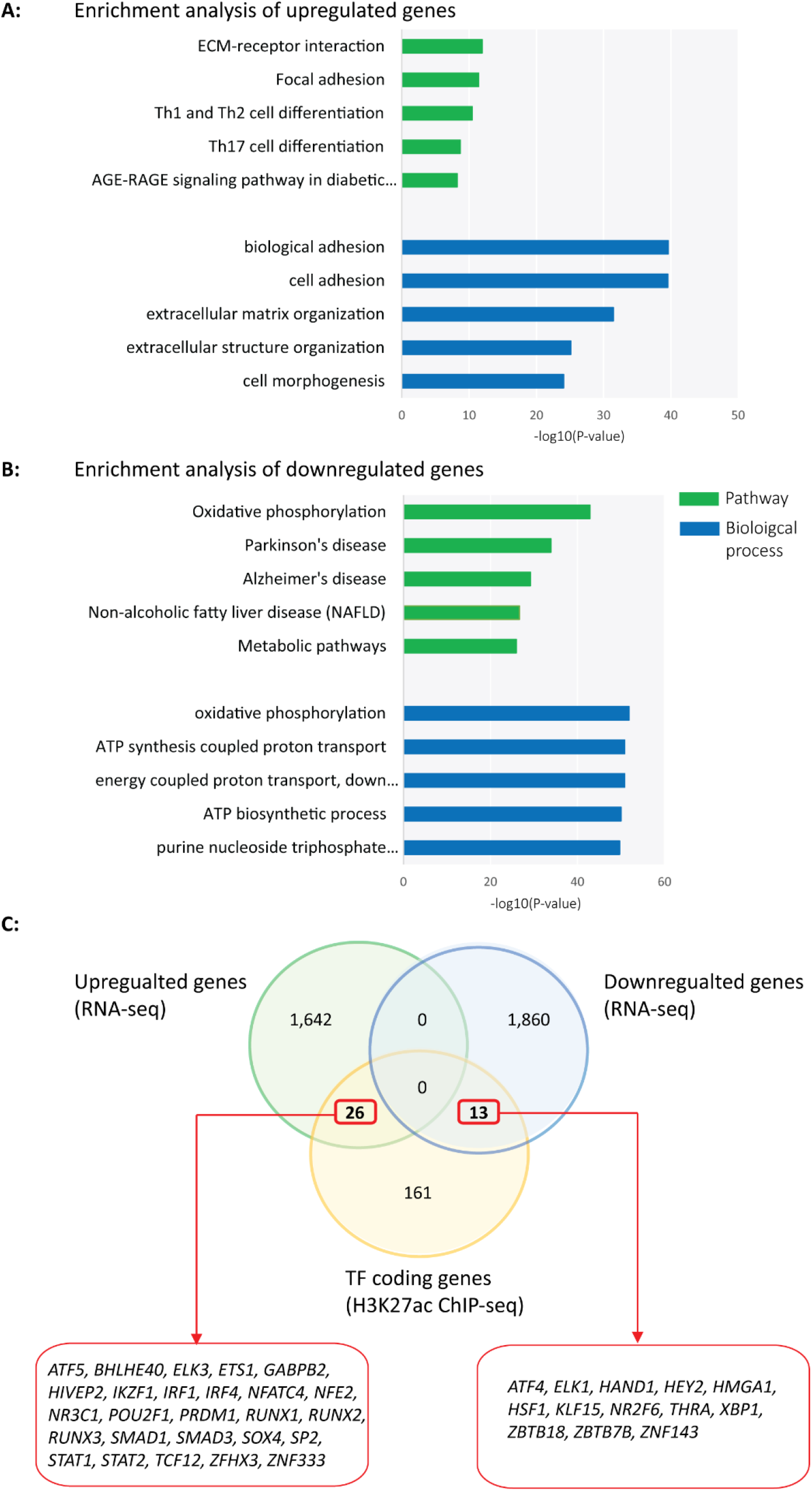
Differentially expressed genes between patients and controls (RNA-seq): **A)** Top 5 enriched Pathway and GO: Biological process terms by upregulated genes in PLN versus control hearts. Notably, the most upregulated genes are part of fibrosis pathways and extracellular matrix organization. **B)** Top 5 enriched Pathway and GO: Biological process terms by downregulated genes in PLN versus control hearts. Note the enrichment of metabolic pathways. **C)** Transcription factor (TF) coding genes, which were predicted to bind to enriched motifs in H3K27ac ChIPseq differentially acetylated regions, with altered mRNA expression levels between PLN and control hearts.

### Integrative analysis of the ChIP-seq and RNA-seq data in PLN versus control hearts

To investigate the upstream players that could play a major role in regulating the subsequent biological processes, we performed an integrative analysis using both the ChIP-seq and the RNA-seq data. We collected all 200 TF-encoding genes, which were annotated from the enriched TFBMs in differentially acetylated regions using the ChIP-seq data, and examined their mRNA expression changes between the PLN and the control groups using the RNA-seq data. As a result, 26 and 13 annotated TF coding genes also showed significantly up- and downregulated mRNA expression levels in PLN versus control hearts (**Figure 6C**). Notably, several overlapping TF coding genes are well-known to play a critical role in the most enriched GO terms and pathways using the ChIP-seq and/or RNA-seq data, such as ECM-related *SMAD3* (upregulated) and metabolism-related *KLF15* (downregulated).

### Validation of accumulated lipid droplets and impaired fatty acid oxidation (FAO) in PLN versus control hearts

Next, we performed histological characterization of the control and PLN hearts. We performed Oil Red O (ORO) staining and observed that lipid droplets were more accumulated in cardiomyocytes of the PLN hearts than the controls, suggesting an insufficient FAO (**Figure 7A**). Besides, we performed schematic digitalized quantification of the percentage of fibrosis, adipocytes, and cardiomyocytes based on Masson’s trichrome staining based on a technique we established previously^8,10^, We see a striking biventricular fibrofatty tissue replacement (myocardial adipogenesis) mostly in the subepicardial region of the myocardium (**Figure 7B**). Analysis of the differentially expressed genes in the FAO pathway shows that 25 out of 30 genes are downregulated in PLN vs. controls (**Figure 7C**). Furthermore, we performed transmission electron microscopy on control and PLN hearts and observed increased lipid droplet formation (significantly higher LD area, diameter, and aspect ratio) and disrupted integrity of mitochondria, the key organelle where FAO takes place (**Figure 8**). Combined, these findings are in line with our omics data indicating the impaired FAO in PLN hearts.

**Figure 7:**
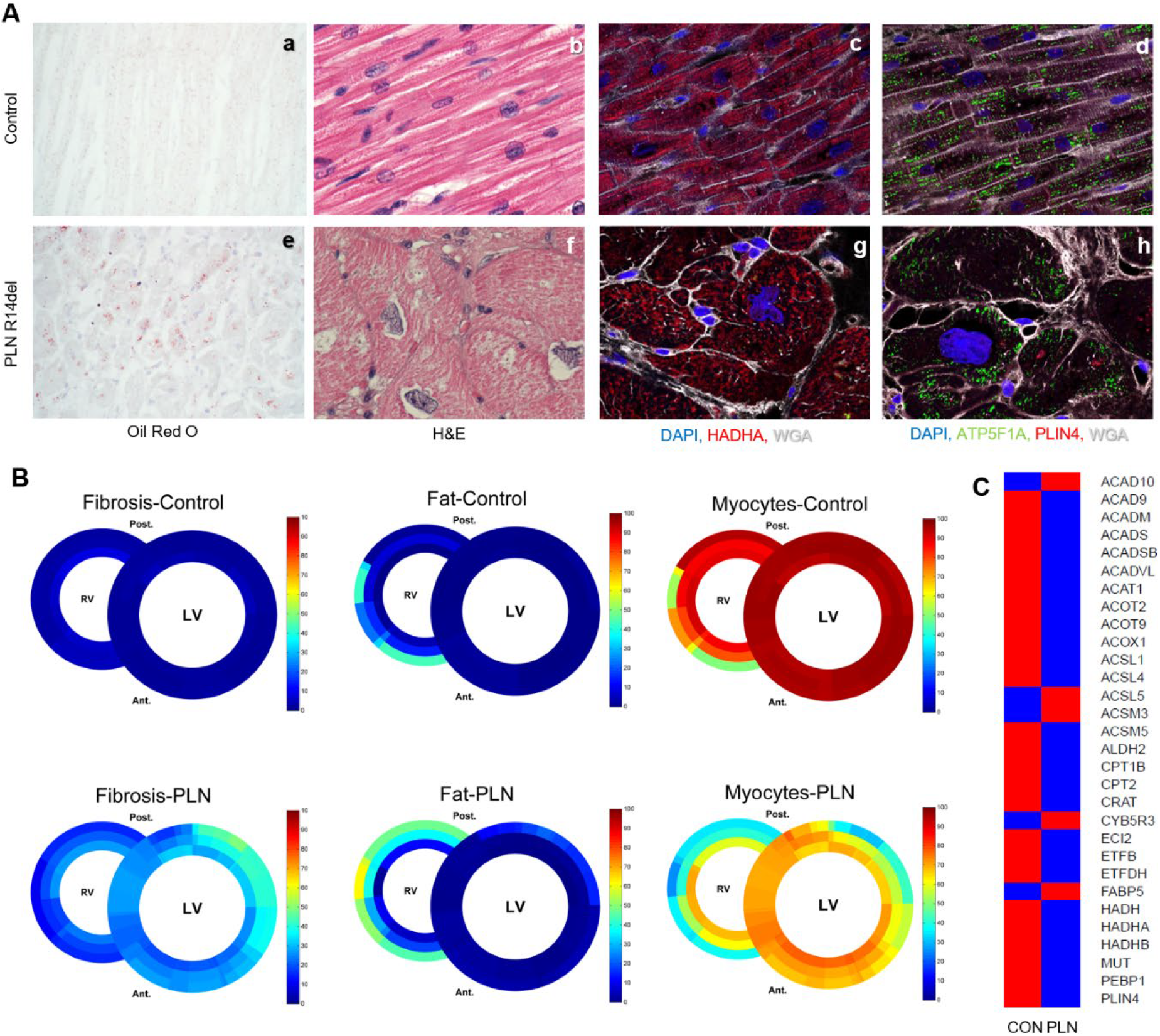
Validation of impaired (lipid) metabolism in PLN hearts. **A)** Microscopic analysis control and PLN R14 del cardiac tissue. **a,e**: lipid droplet formation based on Oil Red O (ORO) staining showing increased lipid droplets accumulation in PLN hearts; 40x magnification; **b,f:** hematoxylin-eosin (H&E) staining; 63x magnification; **c,g**+**d,h**: immunofluorescence analysis of DAPI (nuclei), WGA (wheat germ agglutinin for cell membranes), HADHA and ATP5F1A (mitochondria), and PLIN4 (lipid droplets); 63x magnification. Note the parallel cardiomyocyte orientation with normal-shaped nuclei in controls while hypertrophic misshaped cardiomyocytes with abnormal nuclei in PLN. **B)**. Schematic digital representation of the percentage of fibrosis, adipocyte infiltration, and cardiomyocyte distribution between control and PLN group. Imagine analysis if based on Masson’s trichrome. **C)** Differential expression of genes involved in fatty acid oxidation pathway based on RNA-seq analysis between control and PLN hearts. Average per group is indicated, red - upregulated, blue - downregulated.

**Figure 8:**
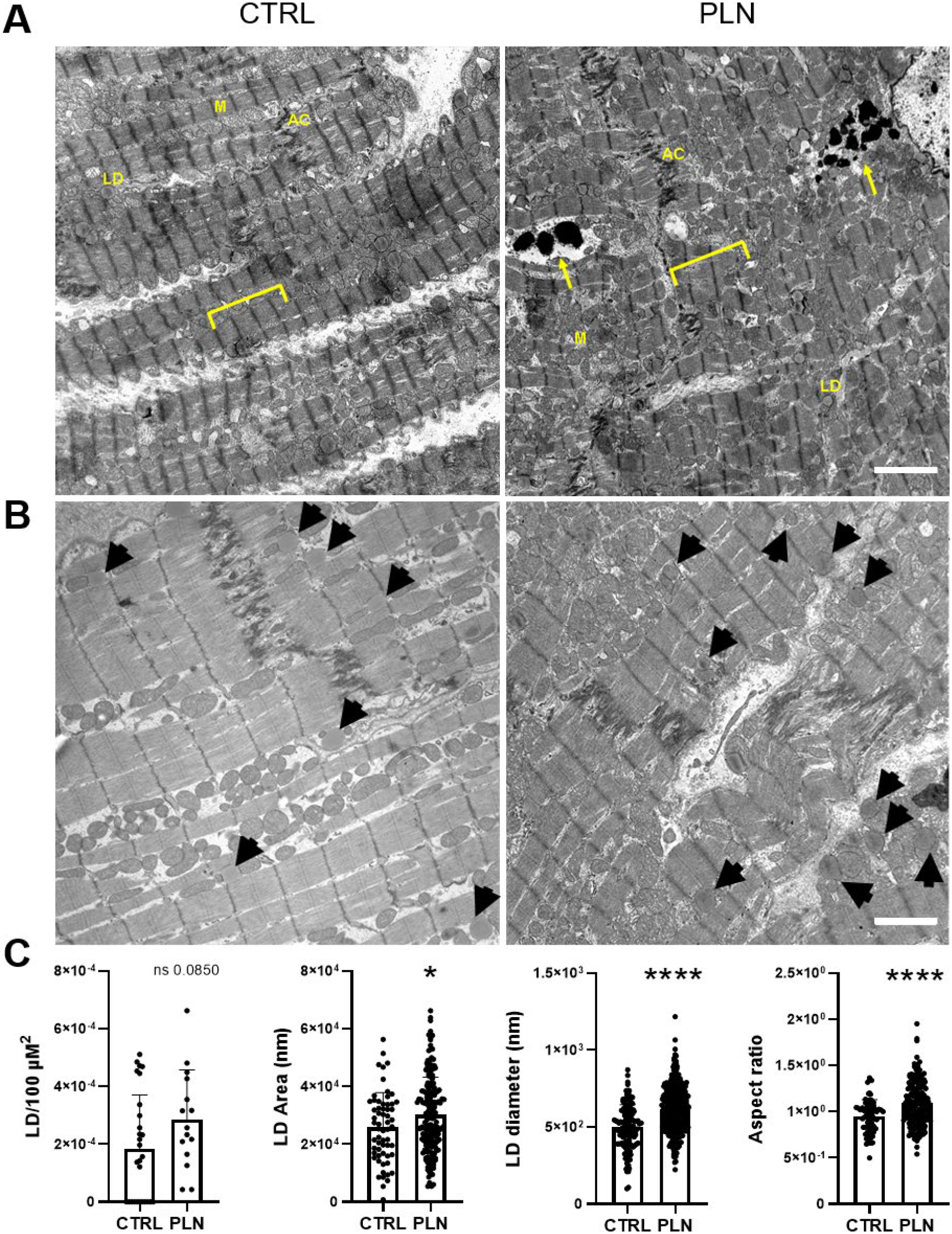
Ultrastructural characterization of PLN R14del hearts. **A)** Left: Transmission electron microscopy image of heart tissue from a healthy individual showing regular myofilaments arranged in sarcomeres (brackets), variable-sized mitochondria (M) arranged in rows in between myofibrils, lipid droplets (L) and profiles of the intercalated disc with a large mixed-type junction (area composita, AC) in between cardiomyocytes. Right: Transmission electron microscopy image of heart tissue from PLN R14del mutation-carrier (PLN) showing less regular myofilaments arranged in sarcomeres (brackets), variable-sized mitochondria (M) irregularly located around the myofibrils, lipid droplets (L) and profiles of the intercalated disc with a large mixed-type junction (area composita, AC) in between cardiomyocytes. Note the presence of structurally abnormal protein aggregates in black (arrows). **B)** Left: Transmission electron microscopy image of heart tissue from a healthy individual showing lipid droplets (black arrows). Right: Transmission electron microscopy image of heart tissue from p.ArgR14 mutation-carrier (PLN) showing lipid droplets (black arrows. **C)** Multiple shape descriptors were determined for manually traced lipid droplets. Abbreviations LD; lipid droplets. n = 3 healthy individuals, 67 LD, n=2 PLN individuals, 177 LD). Data are expressed as mean ±SD, *p<0.05, ****p<0.0001, unpaired T-test was used to compute statistical significance. Scale bar: 2 μM.

## DISCUSSION

This study provides unprecedented information on changes of chromatin activity and global transcriptional regulation in the end-stage hearts from cardiomyopathy patients carrying PLN R14del mutation when compared with the healthy donor hearts. Integrating these multi-omics data in a hypothesis-free manner, we showed and disturbed lipid metabolism in PLN hearts and identified several key transcription factors (e.g. *PPARA* and *KLF15*), which played a critical role in regulating the altered metabolism. It has been shown that regulatory regions display a high level of tissue specificity and that enrichment of GO terms based on the annotation to genes with the closest TSS has cell-specific patterns.^38,39^ Here, we confirm that information based on chromatin activity in heart tissue from both annotated genes in the vicinity of differentially acetylated regions as well as their TFBM enrichment point to known pathways involved in end-stage cardiomyopathies, such as fibrosis, cell death, or impairment of muscle function and lipid (energy substrate) metabolism.^40,41,42^

We have annotated several notable genes to differentially acetylated regions. For example, the regulatory region in the vicinity of FST had the strongest observed hyperacetylation status in our dataset. It is associated with extracellular matrix-related and calcium-binding proteins and its expression is altered during heart failure.^43^ Differentially acetylated regions were detected also within ±5kb from TSS of *CHKB, DES, FHL2, HADHA/HADHB, KCNQ1, MLYCD, MYH7, MYL3, PDLIM3, PLEC*, and *PNPLA2*, and binding motifs were enriched for TFs *GATA4* and *POU1F1*, genes previously were shown to cause genetic forms of dilated cardiomyopathy.^44^ It has to be noted we have not detected a significant difference between the level of H3K27ac binding in the *PLN* gene itself, nor its binding partners such as *ATP2A1, ATP2A2, PRKG1, PRKG2,* or *EDA* (Source: Gene Cards, STRING).

By comparing with diseased hearts without PLN R14del-related cardiomyopathies, we identified a list of PLN R14del-specific regions and annotated several notable genes in these regions, including many known genes involved in lipid metabolism (e.g. *AGPAT2, GLIPR1, HADHA/HADHB, MLYCD, MSMO1, PLPP1, PTGDS,* and *INPP5J21*). In addition, notable extracellular matrix-associated genes involved in fibrosis formation have been found, such as *COL6A3, COL8A1, COL14A1, CFT1, LAMB2, LUM,* and *MMP223,* or genes crucial for sarcomeric contraction: *CNN3, DES, MB, MYH7, MYL3, MYL7*, and *MYLPF19.*

Besides, we also identified differentially expressed genes in PLN versus control hearts using RNA-seq and showed similar enrichment results, including the activated fibrosis formation and the suppressed energy metabolism. Notably, many annotated genes from PLN R14del-specific regions also showed significantly differentially expressed mRNA expression between PLN and control hearts, such as *HADHA, HADHB, PTGDS, COL14A1,* and *LUM*.

By integrating H3K27ac ChIP-seq and RNA-seq data, we identified 39 TF coding genes that could play a critical role in regulating the pathological mechanisms in PLN R14del hearts. *KLF15*, one identified TF coding gene from the integrative analysis with a suppressed mRNA level, is a key regulator in cardiac lipid metabolism.^45^ A decreased FAO and increased glycolysis are observed in *KLF15*-insufficient hearts^46^. Furthermore, the direct targets and the direct cooperator of *KFL15* such as *PDK4* and *PPARA*, in the regulation of FAO have also been shown in our H3K27ac ChIP-seq and/or RNA-seq data.

Despite these well-characterized metabolic disturbances in diseased hearts, the metabolic impairment remains difficult to study in PLN R14del cardiomyopathy. A previous study using the murine model demonstrated that DCM hearts carrying another mutation (p.Arg9Cys) in PLN showed increasing mRNA levels of profibrotic cytokines and ECM markers from the early to the severe stages, such as TGFβ, connective tissue growth factor, periostin, and thrombospondin.^47^ Notably, transcriptional changes in cardiomyocytes from the early stage DCM hearts showed significant repression of genes in aerobic metabolism (e.g. FAO and oxidative phosphorylation) and activation of genes in glucose metabolism when compared with the wild type cardiomyocytes, indicating the switch from fatty acid to glucose utilization occurs at the early stage. Furthermore, by comparing cardiomyocytes from the end-stage DCM hearts and controls, PPAR signalling was the most enriched metabolic pathway by downregulated genes, including Ppara, Ppargc1a, Rxrg, Klf15.

Although fat is a normal component of the heart, especially at the epicardium,^48^ the precise pathological mechanism underlying fat infiltration related cardiomyopathies remains to be elucidated.^49^ A previous study showed the possibility of differentiating adipose-derived stromal vascular fraction cells into cardiomyocyte-like cells.^50^ Another study showed the transduction of cardiac mesenchymal progenitors into adipocytes.^51^ Additionally, fibroblasts can also be differentiated into adipocytes, directly and indirectly, promote the differentiation of adipocytes indirectly via the adipose progenitor cells.^52,53^ These studies indicate the complex networks between cardiomyocytes and non-myocyte cell types during the metabolic (mal)adaption. Therefore, future studies are needed to investigate the cell-to-cell interactions during the cardiac energy rearrangement in PLN R14del cardiomyopathy.

Taken together, we have catalogued valuable information of the chromatin acetylation activities and transcriptome regulations in PLN R14del hearts when compared with controls. Combining these data, we obtained the disturbed energy metabolism and identified several key metabolic regulators, which were further confirmed using multiple functional assays. Due to the difficulties in obtaining cardiac samples from the early stage PLN R14del hearts and the lack of human mimicking clinical phenotypes in the PLN R14del animal models,^1^ the in-depth analysis of integrating data using *ex vivo* end-stage PLN hearts presented in this study revealed for the first time the association between PLN R14del mutation, the impairment of FAO, and the intracellular lipid accumulation. Additionally, identified TFs and genes involved in the metabolic pathway could contribute to the pathological progression of PLN R14del cardiomyopathy, and they serve as promising therapeutic targets to further treatments.

## ACKNOWLEDGMENTS

We are very thankful to the patients and their families for providing valuable cardiac tissue for research purposes. We further thank Joyce van Kuik and Erica and Sierra-de Koning for technical assistance with tissue processing and ChIP-seq library preparation.

## SUPPLEMENTARY FILES

**Table S1:** Detailed clinical overview of the included individuals.

**Table S2:** An overview of H3K27 ChIP-seq data obtained from cardiac tissues in 4 analyses.

**Table S3:**

A. Differentially acetylated regions between PLN and control hearts.
B. Genes annotation of hyperacetylated H3K27ac ChIPseq regions between PLN and control group using a 5kb TSS window.
C. Genes annotation of hypoacetylated H3K27ac ChIPseq regions between PLN and control group using a 5kb TSS window.

**Table S4:**

A. Gene set enrichment analysis of annotated genes from hyperacetylated regions in PLN versus control hearts.
B. Gene set enrichment analysis of annotated genes from hypoacetylated regions in PLN versus control hearts.

**Table S5:**

A. Enriched transcription factor binding motifs in differentially acetylated regions and TFs coding genes,
B. Gene set enrichment analysis of transcription factor coding genes.

**Table S6:**

A. Altered regulatory regions that were specific in PLN R14del hearts when compared with the controls and hearts with ischemic or non-ischemic dilated cardiomyopathy.
B. Annotated genes from the hyper- and the hypoacetylated regions that were PLN-specific when compared with the other groups.

**Table S7:** Differentially expressed genes between PLN and control hearts.

**Table S8:**

A. Gene set enrichment analysis of upregulated genes in PLN versus control hearts.
B. Gene set enrichment analysis of downregulated genes in PLN versus control hearts

